# A unifying framework for summary statistic imputation

**DOI:** 10.1101/292664

**Authors:** Yue Wu, Eleazar Eskin, Sriram Sankararaman

**Affiliations:** Department of Computer Science, UCLA; Department of Human Genetics, UCLA

## Abstract

Imputation has been widely utilized to aid and interpret the results of Genome-Wide Association Studies(GWAS). Imputation can increase the power to identify associations when the causal variant was not directly observed or typed in the GWAS. There are two broad classes of methods for imputation. The first class imputes the genotypes at the untyped variants given the genotypes at the typed variants and then performs a statistical test of association at the imputed variants. The second class of methods, summary statistic imputation, directly imputes the association statics at the untyped variants given the association statistics observed at the typed variants. This second class of methods is appealing as it tends to be computationally efficient while only requiring the summary statistics from a study while the former class requires access to individual-level data that can be difficult to obtain. The statistical properties of these two classes of imputation methods have not been fully understood. In this paper, we show that the two classes of imputation methods are equivalent, *i.e.*, have identical asymptotic multivariate normal distributions with zero mean and minor variations in the covariance matrix, under some reasonable assumptions. Using this equivalence, we can understand the effect of imputation methods on power. We show that a commonly employed modification of summary statistic imputation that we term summary statistic imputation with variance re-weighting generally leads to a loss in power. On the other hand, our proposed method, summary statistic imputation without performing variance re-weighting, fully accounts for imputation uncertainty while achieving better power.

## Introduction

Genome-Wide Association Studies(GWAS) has been successfully used to discover genetic variants, typically single nucleotide polymorphisms (SNPs), that affect the trait of interest [1–7]. GWAS measure or type the genotypes of individuals at a chosen set of SNPs and, then, perform a statistical test of association between a given SNP and the trait of interest. SNPs at which the null hypothesis of no association between the genotype and the trait can be rejected are said to be associated with the trait. The threshold that the absolute value of association statistics pass to reject null hypothesis is also referred as significance level.

In a typical GWAS, due to the cost considerations, only a subset of SNPs are genotyped (typed SNPs). Thus, a direct analyses of typed SNPs is likely to have reduced power to detect associations between untyped SNPs and the trait. Thus, imputation methods, that aim to fill in “data” at the untyped SNPs, have emerged as a powerful strategy to increase the power of GWAS. These methods all rely on the correlation or linkage disequilibrium (LD) [8, 9]. between genotypes at untyped SNPs and those at typed SNPs [10–16] Initial work on imputation focused on the problem of genotype imputation, *i.e.*, inferring the genotypes at untyped SNPs given the genotypes at typed SNPs. Genotype imputation methods rely a reference panel in which individuals are typed at all SNPs of interest to learn the LD patterns across SNPs. Given a target dataset in which genotypes are typed at a subset of the SNPs, these methods rely on the LD patterns learned from the reference panel to infer the genotypes at the remaining untyped SNPs.

In the context of GWAS, there are two broad classes of imputation methods to estimate the association statistics at untyped SNPs. The first class relies on genotype imputation to infer the genotypes at the untyped SNPs followed by computing association statistics at the imputed genotypes [10–14, 16]. We refer to this class of imputation methods as **Two-step imputation** methods. In practice, the most successful methods for the first step of genotype imputation are based on discrete Hidden Markov Models (HMM) [10, 16]. The second class of methods directly imputes the association statistics at the untyped SNPs given the association statistics at the typed SNPs. As shown in previous work [17, 18], the joint distribution of marginal statistics at the typed SNPs and untyped SNPs follow a multivariate normal distribution (MVN) [17–21]. This class of methods utilizes the correlation between the association statistics induced by their dependence on the underlying genotypes [22, 23]. This class of methods is termed *summary statistic imputation* (**SSI**). Summary statistic imputation is appealing as it tends to be computationally efficient while only requiring the summary statistics from a study while the first class requires access to individual-level data which can be difficult to obtain in practice. Current summary-statistic based imputation methods calibrate the imputed statistics using a technique we call *variance re-weighting* (**SSI-VR**). Despite recent progress, the statistical properties of summary statistic imputation methods (including the impact of variance re-weighting) and the connection between the two classes of summary statistic imputation methods has not been adequately understood.

In this paper, we show that the two classes of imputation methods, **Two-step imputation** and **SSI** are asymptotically multivariate normal with small differences in the underlying covariance matrix. Using this asymptotic equivalence, we can understand the effect of the imputation method on power. Our new method, SSI, perfoms summary statistic imputation without variance re-weighting. The resulting statistics do not then have unit variance as in traditional summary statistic imputation but instead correctly take into account the ambiguity of the imputation process.

We compared the peroformance of the imputations methods on the Northern Finland Birth Cohort (NFBC) data set [24] to show that SSI increases power over no imputation while SSI-VR can sometimes lead to lower power. Finally, we ran SSI, SSI-VR and Two-step imputation on the NFBC dataset and show that the resulting statistics are close thereby justifying the theory.

## 2 Results

### 2.1 Overview of Summary Statistics

Assume we have a total of *M* = (*U* + *O*) SNPs that are partitioned into *O* observed (or tag) SNPs {*snp*_1_, *snp*_2_, *snp*_3_…*snp*_*O*_} and *U* missing SNPs {*snp*_1_, *snp*_2_, *snp*_3_,…*snp*_*U*_} for *N* individuals. For the *O* tag SNPs, let ***s***_*O*_ be a vector of association statistics of length *O*, ***λ***_*O*_ be a vector of non-centrality (NCP) parameters of length *O*, and let **Σ**_*O*_ be a *O* × *O* matrix of their pairwise correlation coefficients. For the *U* missing SNPs, let ***s***_*U*_ be a vector of association statistics of length *U*, ***λ***_*U*_ be a vector of NCP parameters also of length *U*, and let **Σ**_*U*_ be a *U* × *U* matrix of their pairwise correlation coefficients.

Let **Σ**_*UO*_ be a *U* × *O* matrix of the pairwise correlation, *i.e.*, linkage disequilibrium (LD), between missing SNPs and observed SNPs. Thus, we have a *M* × *M* LD matrix, **Σ**_*LD*_. We can partition the LD matrix as: 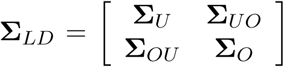. For large sample sizes, the association statistics follow a multivariate normal distribution,

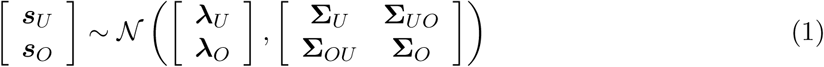

Under the null where we assume that none of the SNPs is causal, ***λ***_*U*_ and ***λ***_*O*_ are equal to **0**.

### 2.2 Example

**Figure 1:**
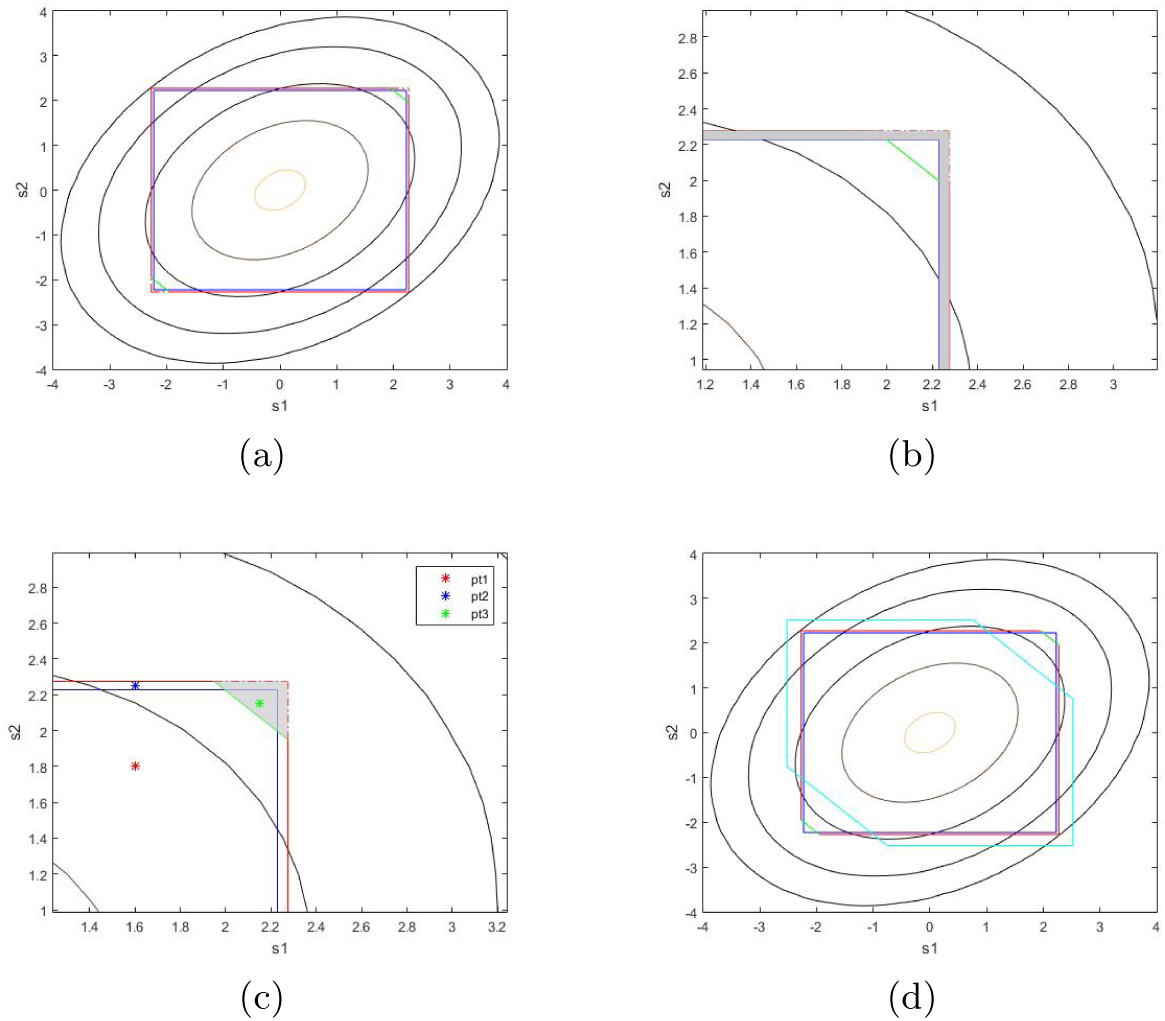
The effect of imputation on the rejection boundary. This figure shows rejection boundary with no imputation, with imputation (SSI), and variance re-weighted imputation (SSI-VR) for an example containing two observed SNPs *snp*_1_, *snp*_2_ and an unobserved SNP *snp*_3_. The contours represent the probability density of the statistics for the observed SNPs: *s*_1_ and *s*_2_ projected in the plane. In Figure 1a, the blue box is the rejection boundary with FWER 0.05 for *snp*_1_ and *snp*_2_ before imputation. The polygon with red and green colored boundaries is the rejection boundary after imputation. Figure 1b and Figure 1c are a zoomed in version of Figure 1a to show the rejection boundaries changes. Figure 1b shows the power change on two observed SNPs. Figure 1c shows the power change on the imputed SNP and has 3 points corresponding to different scenarios. Figure 1d shows the rejection boundary of imputation with SSI-VRin cyan color in addition to the rejection boundaries seen in Figure 1a.

We consider a simple example to illustrate how imputation affects the rejection threshold at a given set of SNPs. We consider three SNPs: *snp*_1_, *snp*_2_, and *snp*_3_. In this example, *snp*_1_*, snp*_2_are observed, and *snp*3 is imputed. We assume the statistics of the tag SNPs (*snp*_1_*, snp*_2_), 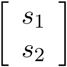 follows 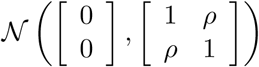 where |*ρ*| ≤ 1 and we use π(*s*_1_, *s*_2_) to denote this distribution. We also assume that the statistics of the tag SNPs *snp*_1_, *snp*_2_ and the unobserved SNP *snp*_3_ jointly follow the distribution 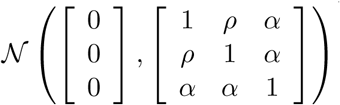 where |*ρ*| ≤ 1, |*α*| ≤ 1.

Thus having the joint distribution of the statistics *s*_1_, *s*_2_, and *s*_3_, we can compute the conditional distribution of the untyped SNP conditioned on the marginal statistics of the typed SNPs *s*_1_ and *s*_2_:

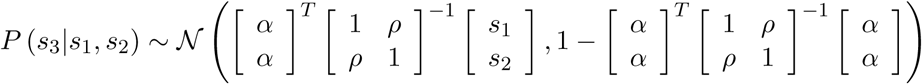

Typically, summary statistic imputation uses the posterior mean of the statistic *s*_3_ given the observed values of 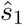 and 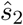 to estimate *s*_3_. In our example, this leads to the statistic *s*_3_ for *snp*_3_ being imputed as a function of 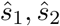:

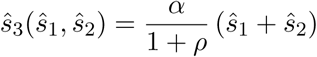

We choose thresholds *t* for rejecting each of the statistics 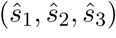 such that the family-wise error rate, *i.e.*, the probability of at least one false positive, is controlled at a level 0.05. For each tested SNP, we choose the threshold to be the same.

In the case where no imputation is performed, we only test two SNPs. We use the same threshold *t* for SNPs *snp*_1_ and *snp*_2_. Figure 1a shows the rejection boundary (the blue box) for two SNPs with correlation *ρ* = 0.36 where the region outside this box corresponding to the rejection region. Given the joint density *π*(*s*_1_*, s*_2_) of the association statistics (*s*_1_*, s*_2_), we determined the rejection boundary by computing the length of the side of the blue box such that the cumulative density in the rejection area, *i.e.*, the area under the density *π*(*s*_1_*, s*_2_) outside the box is equal to 0.05. Mathematically, we need to find *t* such that *F W ER*(*t*) = 0.05 where:

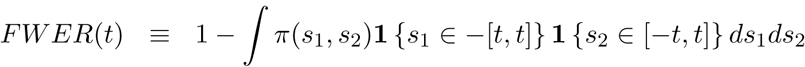

Here **1** {*s*_1_ ∈ − [*t, t*] **1** {*s*_2_ ∈ − [*t, t*] defines the acceptance region, *i.e.*, the set of points (*s*_1_, *s*_2_) ∈ ℝ^2^ where the null hypothesis at both SNPs are accepted.

We now consider the effect of testing imputed SNPs in addition to the tag SNPs. The rejection region for *snp*_1_, *snp*_2_, *snp*_3_ are the regions outside the intervals *R*_1_ = [−*t*, *t*]*, R*_2_ = [−*t*, *t*], *R*_3_ = [−*t*, *t*] respectively. We can compute the FWER for a given *t* by determining the probability mass outside the rejection region. To do this, we note that the joint sampling distribution of 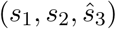 is determined only by the distribution of (*s*_1_, *s*_2_) since 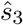 is a deterministic function of *s*_1_ and *s*_2_.

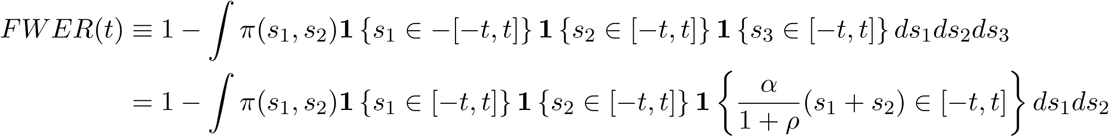

Notice that, in the setting with imputation, the acceptance region **1** {*s*_1_ ∈ [−*t*, *t*]}**1** {*s*_2_ ∈ [−*t*, *t*]} 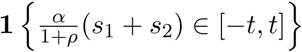 can never increase relative to the setting where only the tag SNPs are tested. Now consider the case where the null hypothesis at both the observed SNPs is accepted. This happens when 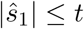 and 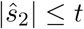. Then the statistic at the imputed SNP:

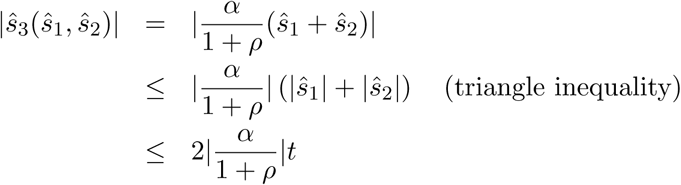

Thus, if 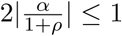, then we have 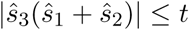. Thus, the imputed SNP will never be rejected when neither of the observed SNPs is rejected. Thus, the acceptance region remains the same as the setting when only the tag SNPs are tested. In other words, imputation does not change the rejection boundary.

On the other hand, when 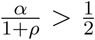, then imputation will change the rejection region. Figure 1 shows the effect of imputation with *α* = 0.80 and *ρ* = 0.36 so that 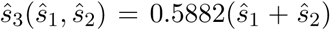. The rejection boundary of the observed SNPs *snp*_1_ and *snp*_2_ after imputation are shown by the red lines. The rejection region for *snp*_3_ corresponds to the region where |0.5882(*s*_1_ + *s*_2_)| > *t* which corresponds to the green line. Thus the cumulative density outside the polygon of red and green lines is the same as the rejection area outside the blue box. In Figure 1b, the shaded area indicates the power loss on the observed SNPs, and in Figure 1c the shaded area is the power gained from imputation.

Thus assume we have three points, *p*1, *p*2 and *p*3 in Figure 1c, which are three different pairs of association statistics of observed SNPs *snp*1 and *snp*2. The first point is in both the blue rectangle and the polygon, which means we will accept null with or without imputation. The second point *p*2 is the case that, without imputation we will reject null, and after imputation we will accept null because of the change of boundary on observed SNPs. The third point *p*3 is the special case. In this case, the observed SNPs don’t have significant association because it lies inside the blue box, but after imputation, the imputed SNP has a significant association since it lies outside the polygon and thus we reject the null.

### 2.3 Simulation Results

As shown in previous work on summary statistics [22], the marginal statistics at typed SNPs and untyped SNPs follow a multivariate normal distribution. With the assumption that none of the SNP is significantly associated with train, the mean of the multivariate normal distribution is 0.

As in the previous simple case having 3 SNPs, *snp*_1_*, snp*_2_ and *snp*_3_, under the null hypothesis of no association, the summary statistics follow the distribution 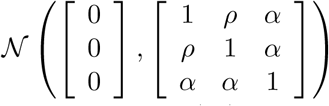.

Thus having the joint distribution of the statistics *s*_1_, *s*_2_, and *s*_3_, we can compute the conditional distribution of the untyped SNP conditioned on the marginal statistics of the typed SNPs *s*_1_ and *s*_2_:

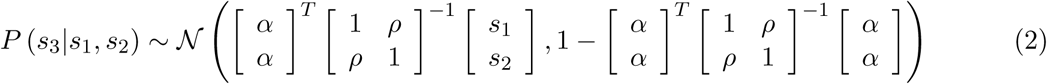

Summary statistic imputation estimates *s*_3_ using the mean of the above distribution 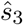. The variance of the imputed statistic: *var*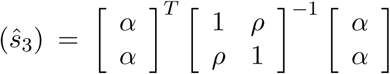 is smaller than 1 (since Equation 2 shows that the variance of is 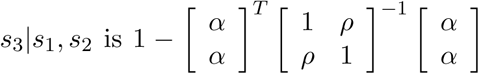 and the variance is non-negative). Thus, in most summary statistics imputation [22, 23], *snp*_3_ is imputed as 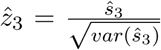 so that all the association statistics have variance 1. Since the variance of 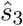 is ≤ 1, the new statistic 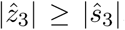. As a result, for a given threshold, the acceptance region in SSI-VR is never greater than with SSI. In other words, to achieve a given FWER, the threshold *t* needs to be larger for SSI-VR than without as shown in Figure 1d.

Now having *snp*_3_ imputed using summary statistics, we want to find out how power is affected by SSI and SSI-VR. In section 3 of the Supplementary Information, we analytically compute the average marginal power function for both methods. In order to assess power, we assume that 3 SNPs, *snp*_1_*, snp*_2_ and *snp*_3_ are drawn from a region associated with a trait. We assume that the untagged variant, *snp*_3_, is causal with non-centrality parameter (NCP) so that (*s*_1_*, s*_2_*, s*_3_) follow a non-zero mean multivariate normal distribution: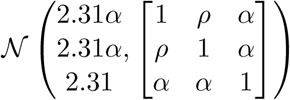. We choose the NCP to be 2.31 so that the maximum power of no imputation will be around 0.5, which will happen when both *α* and *ρ* are 1. We let the correlation between untagged and tag SNPs *α* and the correlation between tag SNPs *ρ* vary across: [0.1,0.2,…, 0.9,1].

For each combination of [*α, ρ*], we determined a set of 3 thresholds i) for no imputation, ii) for imputation, and iii) imputation with variance correction. We drew 10^8^ samples from each distribution, and the power is defined as the the probability that we reject the null hypothesis based on thresholds for each method.

In all the combinations except the cases that the LD matrix is no longer positive definite, we find the power of no imputation, SSI and SSI-VR (Figure 2). In Figure 2a, we compared SSI versus no imputation, and we show that SSI always increases power when 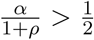 as the ratio is always larger in 1. Since the power of no imputation depend more on the correlation between tagged and untagged SNP, we see the power being sensitive to *α*. For instance, if *α* = 0.7 and *ρ* = 0.3, the average power of no imputation is 0.4918 while the average of the power of imputation with no correction is 0.6614. In figure 2(b), we compared SSI-VR versus no imputation. We see comparing to 2(a), the power increasing much less significant. In fact, in some cases, we observe SSI-VR has less power than no imputation. For example, when *α* = 0.7 and *ρ* = 0.1, the average power of imputation with variance correction is 0.4639, and null has an average power of 0.5154.

**Figure 2:**
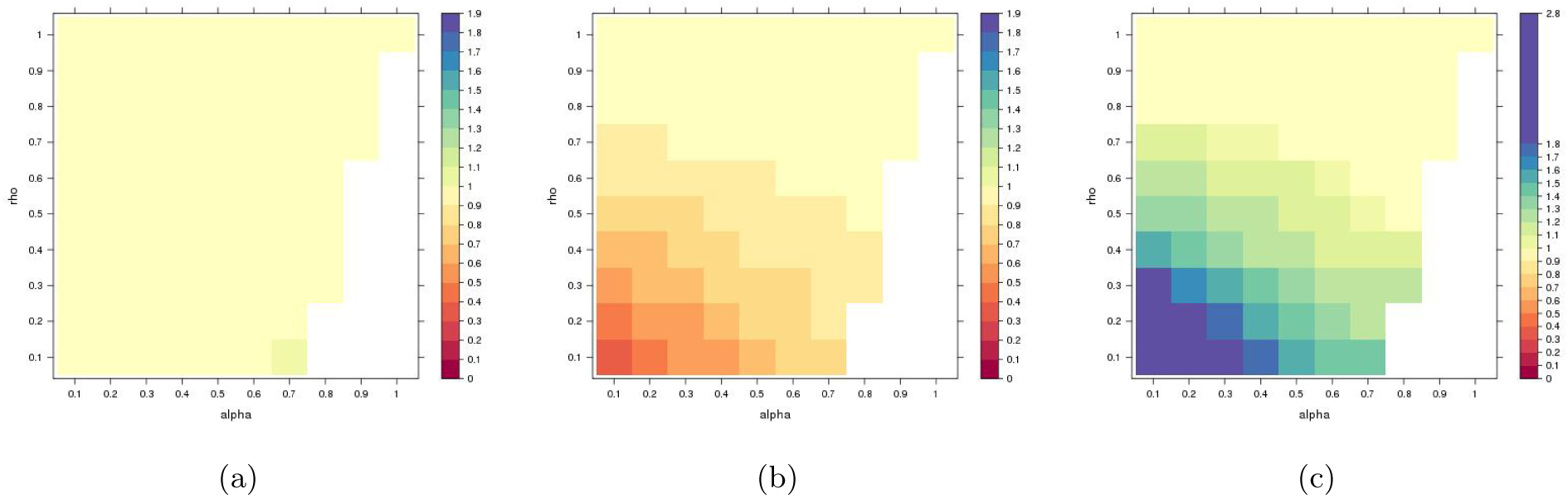
A comparison of the power of imputation (SSI) v.s. no imputation(a), SSI-VR v.s. no imputation(b), and SSI v.s. SSI-VR in a simple example consisting of three SNPs of which only two are observed. In each panel, we plot the ratio of the power of the two methods under all configurations of *α* and *ρ*. In each figure, the configuration of *α* and *ρ* that results in a covariance matrix that is not positive definite, e.g. *α* = 1, *ρ* = 0.1, is left empty. Figure 2a shows that for values of 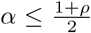, the ratio is near one since the rejection boundary is unchanged (as predicted by our theory) while for values of 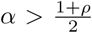, the power of SSI is greater than that of no imputation. Figure 2b and 2c show that SSI-VR can lose power relative to both no imputation as well as SSI for a range of configurations of LD.

Then, we compare imputation and imputation with variance re-weighting in Figure 2c and we notice that SSI-VR will always cause power loss and in figure the value of ratio are all larger than 1. For instance, when *α* = 0.7 and *ρ* = 0.3, the average power of imputation is 0.6614, and the average power of imputation with variance correction is 0.5403.

### 2.4 SSI achieves better power compared to existing methods in Northern Finland Birth Cohort (NFBC)

In order to assess the power of imputation and the effect of SSI-VR on imputation in a real dataset, we simulated marginal statistics utilizing the Northern Finland Birth Cohort (NFBC) dataset.

We assume that every other SNP on chromosome 22 is missing. Thus, we observe half of SNPs on chromosome 22 and perform imputation on the rest. We find the per-SNP threshold for only observed SNPs (*i.e.* no imputation), for SSI and for SSI-VR with the constraint that FWER is controlled at 0.05. We sampled association statics from the multivariate distribution on the observed SNPs from the genome. Then we used the sampled statistics to find the per-SNP significance threshold on the observed SNPs. We found the threshold to be **4.59705**. Having this threshold, we then assume that there are causal SNPs in the genome, *i.e.* the mean of statistics on these SNPs are not 0, and assess the power with no imputation. For no imputation, we found an average power of **0.4946**.

For the imputation methods, SSI and SSI-VR we impute the association statistics using the samples statistics. We impute in two ways, one utilizing the MVN of equation(4), and the other one use variance re-weighting technique as equation(5). Under the null, we found per-SNP thresholds for SSI and SSI-VR to be **4.5977** and **4.6891**. We then assume that there are causal SNPs, and used the thresholds to compute the power of each of the imputation methods. We found the average power to be **0.50124** for SSI and **0.4346** for SSI-VR. Notice that the threshold we found for no imputation, SSI, and SSI-VR are more accurate than Bonferroni correction and thus less conservative.

In the Table 1, we also impute the most significantly associated SNPs reported in previous studies using SSI, SSI-VR and a Two-step imputationusing IMPUTE2 to perform genotype imputation. We find the association statistics are similar across the three methods validating our theoretical results.

**Table 1:** We show that the two classes of imputation method, SSI and Two-step imputation have similar imputation statistics on the NFBC data set. We consider SNPs that were reported significant in a previous study [24]. Then, we treat these SNPs as untyped and impute the marginal statistics using SSI, SSI-VR, and Two-step imputation using IMPUTE2 to impute genotype of untyped SNPs.

| Phenotype | chr | rsID | True Statistics | SSI | \| True - SSI \| | SSI-VR | \| True - SSI-VR \| | IMPUTE2 | \| True - IMPUTE2 \| |
| --- | --- | --- | --- | --- | --- | --- | --- | --- | --- |
| TG | 2 | rs673548 | -5.444 | -5.37 | 0.074 | -5.37 | 0.074 | -4.46 | 0.984 |
|  | 8 | rs10096633 | -5.679 | -5.63 | 0.049 | -5.76 | 0.082 | -5.17 | 0.509 |
|  | 15 | rs2624265 | 4.22 | 3.55 | 0.67 | -3.85 | 0.37 | 3.60 | 0.62 |
| HDL | 15 | rs1532085 | 7.13 | 5.59 | 1.54 | 6.33 | 0.8 | 6.47 | 0.66 |
|  | 16 | rs3764261 | 12.01 | 8.23 | 3.78 | 10.19 | 1.82 | 6.47 | 5.54 |
|  | 16 | rs255049 | 6.06 | 5.11 | 0.95 | 5.5 | 0.56 | 5.70 | 0.36 |
|  | 17 | rs9891572 | 4.25 | 3.99 | 0.26 | 4.02 | 0.23 | 4.40 | 0.15 |
| LDL | 1 | rs646776 | -7.70 | -7.7 | 0 | -7.81 | 0.11 | -6.96 | 0.74 |
|  | 2 | rs693 | 6.81 | 6.27 | 0.54 | 6.34 | 0.47 | 5.91 | 0.9 |
|  | 11 | rs102275 | -4.51 | -4.43 | 0.08 | -4.45 | 0.06 | -4.54 | 0.03 |
|  | 11 | rs174546 | -4.52 | -4.43 | 0.09 | -4.45 | 0.07 | -4.58 | 0.06 |
|  | 11 | rs174556 | -4.69 | -4.73 | 0.04 | -4.85 | 0.16 | -4.62 | 0.07 |
|  | 11 | rs1535 | -4.43 | -4.46 | 0.03 | -4.66 | 0.23 | -4.45 | 0.02 |
|  | 19 | rs11668477 | -5.96 | -3.78 | 2.18 | -4.4 | 1.56 | -5.33 | 0.63 |
|  | 19 | rs157580 | -5.161 | -2.6 | 2.561 | -3.11 | 2.051 | -4.20 | 0.961 |
| CRP | 12 | rs2650000 | -7.08 | -5.25 | 1.83 | -6.54 | 0.54 | -6.05 | 1.03 |
| GLU | 2 | rs560887 | -6.97 | -6.21 | 0.76 | -6.3 | 0.67 | -5.69 | 1.28 |
|  | 7 | rs10244051 | 5.31 | 4.34 | 0.97 | 4.45 | 0.86 | 4.97 | 0.34 |
|  | 7 | rs2191348 | 5.30 | 4.33 | 0.97 | 4.47 | 0.83 | 4.97 | 0.33 |
|  | 11 | rs1447352 | -6.35 | -5.08 | 1.27 | -5.21 | 1.14 | -4.75 | 1.6 |
|  | 11 | rs7121092 | -5.50 | -4.93 | 0.57 | -5.31 | 0.19 | -4.60 | 0.9 |

## 3 Methods

### Summary Statistics

Under the null hypothesis, the joint distribution of the association statistics of the *U* untagged SNP ***s***_*U*_ and the *O* tag SNPs ***s***_*O*_ follows a multivariate normal distribution:

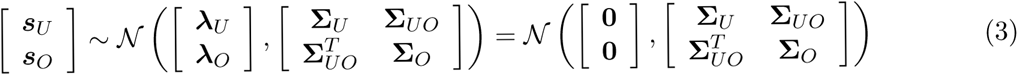

Since none of the *M* = (*U* + *O*) SNPs are associated, the non-centrality parameters of both ***λ***_*U*_ and ***λ***_*O*_ are **0**. Further, the statistics are standardized so that the diagonal elements of the covariance matrix are 1, *i.e.*, Σ_*U*_*i,i*__ = Σ_*O*_*j,j*__ = 1.

#### 3.1.1 Summary statistic imputation

Under the null assumption where ***s***_*O*_ and ***s***_*U*_ are not associated, ***λ***_*U*_ and ***λ***_*O*_ are each **0**. Using the joint distribution, we can compute the distribution of the true statistics at the untagged SNPs, ***s***_*U*_ conditioned on the statistics observed at the tag SNPs, ***s***_*O*_. The conditional distribution follows a multivariate normal distribution, which is computed as follows,

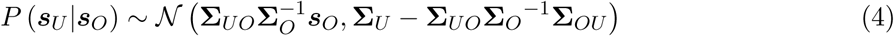

The observed statistics are denoted 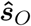. Thus ***s***_*U*_ is imputed using a function of observed statistics:

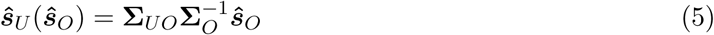

Let ***A*** = **Σ**_*UO*_**Σ**_*O*_^-1^ and thus 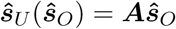.

#### 3.1.2 Summary statistic imputation with variance re-weighting (SSI)

From the previous result, we have 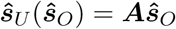. Notice that the underlying joint distribution over the test statistics assumes that each of the statistics at the observed as well as unobserved SNPs has variance one. On the other hand, Equation 5 shows that the variance of the imputed statistic is less than 1. Variance re-weighting proposes standardizing the statistics at the untagged SNPs.

Let *s*_*i*_ be the statistic at the *i*^*th*^ untagged SNP. Thus, instead of imputing *s*_*i*_ using 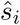, we impute using 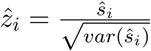, so that all the imputed 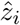 have variance equal to 1. We have: *var*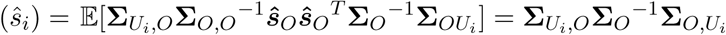. Thus we have:

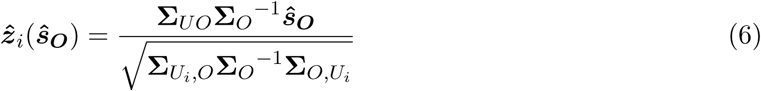

#### 3.2 The impact of imputation on the rejection boundary

SSIuses the following function to impute statistics at the unobserved statistics: 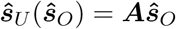. Let ***A***_*i*_ be the *i*^*th*^ row of matrix ***A***, 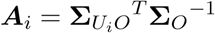, where 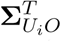 is the correlation vector between untagged variant *snp*_*i*_ and all the observed SNPs. We choose thresholds *t* for rejecting statistics at each of the observed and imputed SNP, *i.e.*, we reject the null hypothesis at observed SNP *O*_*j*_ if 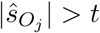 while we reject the null hypothesis at unobserved SNP *U*_*i*_ if 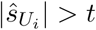 where *t* is chosen to control the FWER. We would like to understand the conditions the threshold *t* for SSIrelative to the threshold *t* when no imputation was performed, *i.e.*, we want to provide conditions when imputation changes the rejection boundary.

##### Theorem 1.

The imputed statistic at snp_i_ computed using SSI will change the rejection boundary iff the sum of the absolute values of all the entries of ***A**_i_*, Σ*_j_*|*A_ij_*| > 1.

*Proof*. See Section 2 in Supplementary Information.

In SSI-VR, instead of using 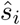 as the imputed statistic for variant *i*, we use

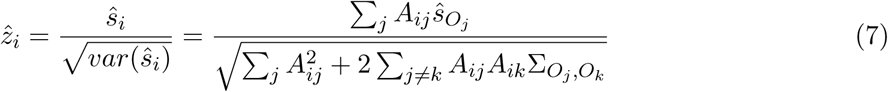

In SSI-VR, untagged variant *i* will effect the rejection boundary iff 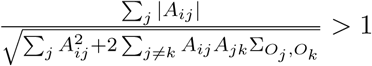.

### 3.3 Two-step imputation

The two-step approach to summary statistic imputation first performs genotype imputation followed by testing for association using the imputed genotypes. Genotype imputation fills in the genotypes at the unobserved SNPs, ***G***_*U*_ given the genotypes at observed SNPs ***G***_*O*_ [15]. Typically, this involves defining a probability distribution for the missing genotypes given the observed genotypes *P* (***G***_*u*_|***G***_*O*_). Let *p*_*i*_(***g***) = *P* (***G***_*U*_*i*__ = ***g***|***G***_*O*_) denote the posterior probability at unobserved SNP *i*. Given a vector ***g*** of *N* genotypes at a SNP, let the association statistic *s*(***g***) be a function of the genotypes ***g***. We can then compute the association statistic at unobserved SNP *i* as the posterior mean of the association statistic: 𝔼 [*s*(***G***_*U*_*i*__)|***G***_*O*_] = Σ_***g***_ ***s***(***g***)*p*_*i*_(***g***). In practice, instead of the posterior mean, association statistics are restricted to imputed SNPs at which the imputation is confident (*e.g.* using the INFO score reported by software such as IMPUTE2 [16]) followed by using the maximum *a posteriori* estimate of the genotype at each SNP. We focus on the posterior mean as it accounts for the uncertainty in imputation and is easier to analyze. We first consider a simple genotype imputation strategy that uses the pairwise correlation among SNPs in a multivariate normal distribution [25] (Section 3.3.1). In Section 3.3.2, we consider the use of hidden Markov Models (HMMs) for genotype imputation.

#### 3.3.1 Genotype imputation using multivariate normal distribution

First, we consider a multivariate normal distribution with mean zero and covariance matrix given by the LD matrix to model the distribution of the genotype vector at the observed and unobserved SNPs for each individual [25]. We can then impute the genotypes for missing SNPs 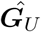 as a function of observed genotypes ***G***_*O*_ using the conditional mean for the multivariate normal distribution (Equation 4). Denoting the *N* × *O* matrix of standardized genotypes as ***X***_*O*_ and the imputed genotype vector across *N* individuals at unobserved SNP *i* as 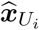, we have:

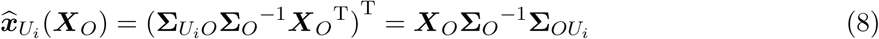

where **Σ**_*U*_*i*__*O*__ is the *i*^*th*^ row of matrix **Σ**_*UO*_.

Given a vector of continuous phenotypes ***y*** ∈ ℝ^*N*^ measured across *N* individuals, the effect size 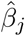 for observed SNP *j* can be estimated by a linear regression of ***y*** on the genotypes at SNP 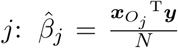 so that the association statistic *s*_*j*_ at this SNP 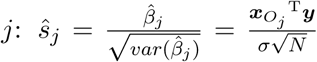. Here *σ* denotes the standard deviation of the phenotype. Analogously, the association statistic 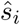 at unobserved SNP *i* is 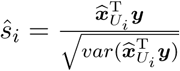. From Equation 8, we have:

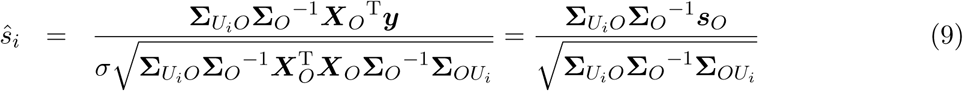

Here we used 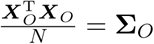.

This function is identical to SSI-VR as seen in Equation 7. Thus, applying the imputation function in Equation 8 to directly impute genotypes is equivalent to SSI-VR.

#### 3.3.2 Genotype imputation using hidden Markov models

We consider the use of a hidden Markov model (HMM) for genotype imputation. These models assume that a reference panel ***M*** is available that contains genotype data across *M* = (*U* + *O*) SNPs [26, 16, 10, 14]. The HMM models the conditional distribution of each of the pair of haplotypes 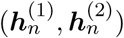 in each of the *N* individuals in the study at the *O* observed and *U* unobserved SNPs by the conditional distribution *P* (***h**|**M***). Specifically, 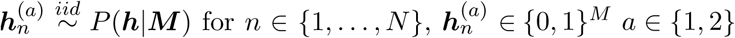.

The effect size estimate for SNP 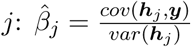 and the association statistic 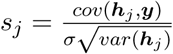.

We show (in Supplementary Information Section 1) that the vector of association statistics asymptotically follows a multivariate normal distribution:

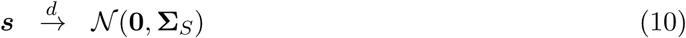

The asymptotic covariance matrix of the association statistics **Σ**_*S*_ depends on the specific HMM used. Under the commonly used Li-Stephens model [27], this covariance matrix is:

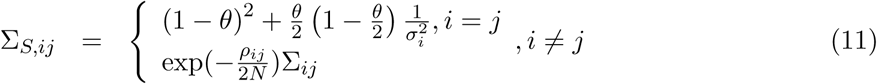

Here Σ_*ij*_ is the LD or the correlation between SNPs *i* and *j*, *θ* is a parameter related to the mutation rate, and *ρ*_*ij*_ is an estimate of the population-scaled recombination rate between SNPs *i* and *j*. Thus, the association statistics computed using genotypes imputed using a HMM follows a multivariate normal distribution with mean zero and covariance matrix equal to a LD matrix with shrinkage applied according to the recombination rate between SNPs.

## 4 Discussion

In this paper, we showed the connection between the two broad classes of imputation, Two-step imputation and SSI. We also showed that a commonly employed modification of SSI, variance re-weighting, will cause power loss using simulation and real data. Thus, this leads us to conclude that SSI (with no variance re-weighting) is more powerful.

Summary statistic imputation assumes that statistics follow multivariate normal distribution: this assumption breaks down for small sample sizes and for rare SNPs. Compared to summary statistics, current HMM methods are likely to be more accurate for rare variation. A possible future direction is to improve accuracy on rare variants and small sample sizes.

## Supplementary Information

### 1 Genotype Imputation using a hidden Markov model

We assume a hidden Markov model for genotype imputation. These models assume that a reference panel ***M*** is available that contains genotype data across (*U* +*O*) SNPs [1–4]. The HMM models the conditional distribution of each of the pair of haplotypes 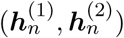 in each of the *N* individuals in the study at the *O* observed and *U* unobserved SNPs by the conditional distribution *P* (***h***|***M***). Specifically, 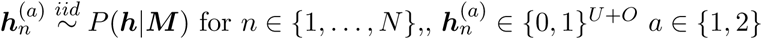.

In GWAS, given a vector of continuous phenotypes ***y*** ∈ ℝ^*N*^measured across *N* individuals, the effect size 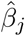 for an observed SNP *j* can be estimated by a linear regression of ***y*** on the genotypes at SNP *j* to obtain an estimate of the effect size. The effect size for SNP *j* is estimated as 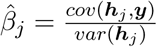 and the association statistic 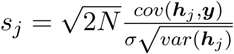.

Let 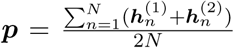. For large sample sizes *N*, ***p*** is asymptotically distributed as a multivariate normal distribution with mean ***µ*** = 𝔼 [***h***|***M***] and covariance matrix 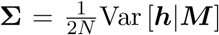. The specific form for the mean and the covariance matrix depends on the form of the HMM used.

For example, in the Li-Stephens HMM [5], [6] showed that

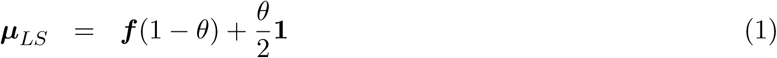

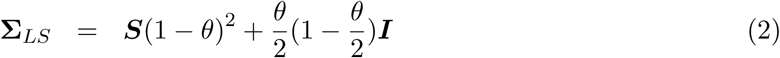

Here ***f*** is the mean allele frequency in a panel, *θ* is a parameter related to the mutation rate, and ***S*** is an estimator of the covariance

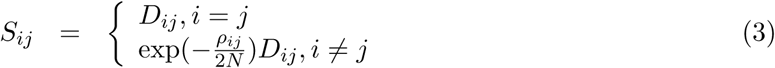

Here *ρ*_*ij*_ is an estimate of the population-scaled recombination rate between SNPs *i* and *j* while *D*_*ij*_ is an estimate of the empirical covariance between SNPs *i* and *j* in a reference panel.

Under the null, we have 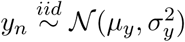 where *y*_*n*_ is independent of the haplotype 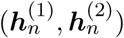.

Let

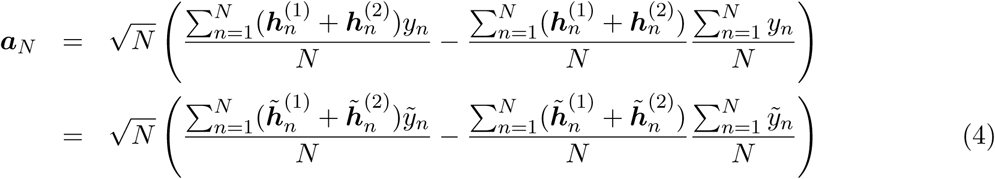

Here 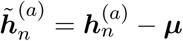 and 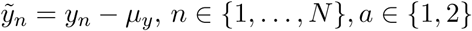.

Now using the Central Limit Theorem and the fact that under the null, the phenotype *y*_*n*_ and the haplotypes 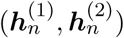 are independent:

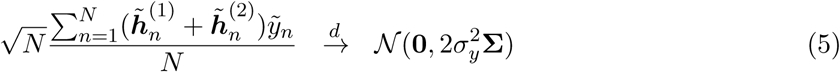

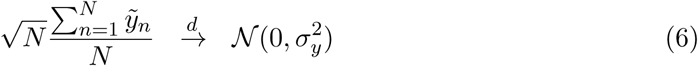

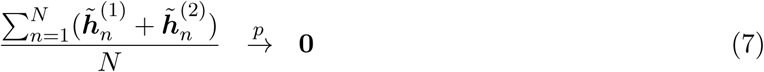

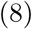

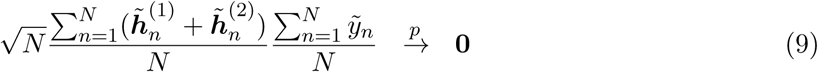

This follows from Equations 6 and 7 by application of Slutsky’s lemma and the continuous mapping theorem [7].

Thus, again applying Slutsky’s lemma to Equations 5 and 9, we can write ***a***_*N*_ as (Equation 4):

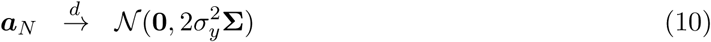

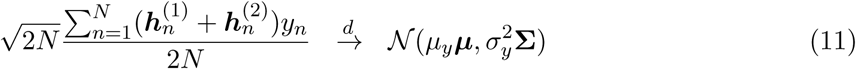

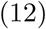

Let 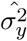 be a consistent estimator of 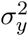. Similarly 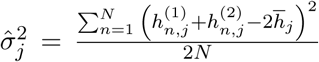 is a consistent estimator of Σ_*jj*_.

Let 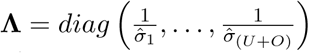 be a diagonal matrix with diagonal entries corresponding to the inverse of the standard deviation of the genotype at each SNP. We can then derive the asymptotic distribution of the asymptotic statistics as:

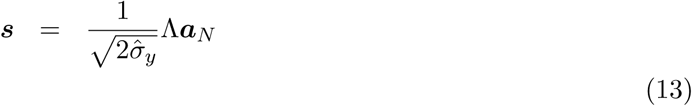

The entry corresponding to SNP *j* is the association statistic for SNP *j*:

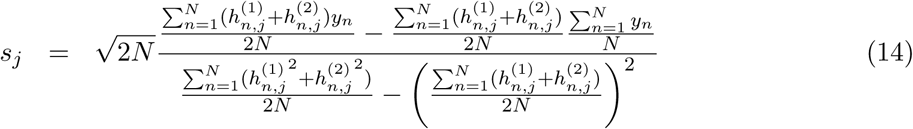

We then have

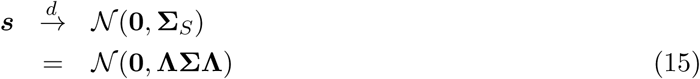

The asymptotic covariance matrix of the association statistics ***s*** under the Li-Stephens model is given by

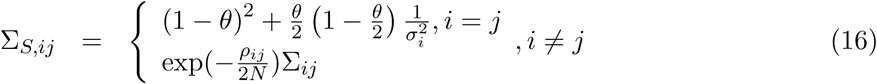

Here Σ_*ij*_ is the LD or the correlation between SNPs *i* and *j*. Thus, the association statistics computed using genotypes imputed using a HMM follows a multivariate normal distribution with mean zero and covariance matrix equal to a LD matrix with shrinkage applied according to the recombination rate between SNPs.

### 2 Impact of summary statistic imputation on the rejection boundary

In general, summary statistic imputation uses a linear function of the observed statistics ***s***_*O*_ to impute the statistic at the unobserved SNP 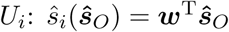 for some weight vector ***w***.

We choose thresholds *t* for rejecting statistics at each of the observed and imputed SNP, *i.e.*, we reject the null hypothesis at observed SNP *O*_*j*_ if |*s*_*O*_*j*__| > *t* while we reject the null hypothesis at unobserved SNP *U*_*i*_ if |*s*_*U*_*i*__| > *t* where *t* is chosen to control the FWER. The acceptance region (the region where all statistics are accepted so that there are no false positives) is the vector of *O* values of the observed statistics that satisfies the following constraints:

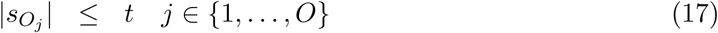

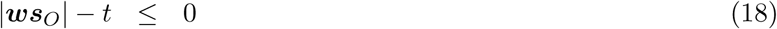

Equations 17 together define a *O*-dimensional hyper-cube with vertices defined by (±*t*,…, ±*t*) while equation 18 defines two hyperplanes ***w***^T^***s**_O_ −t* = 0 and ***w***^T^***s**_O_*+ *t* = 0.

#### Theorem 1.

Unobserved SNP *U_i_* that is imputed using the linear function 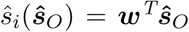 will alter the rejection boundary iff Σ*_j_*|*w_j_*| > 1.

*Proof*. For an untagged variant *snp*_*i*_ to have an effect on the rejection boundary, the two hyperplanes that define the imputed statistic at SNP *i*: |**w**^T^***s**_O_*| -*t* ≤ 0 must intersect the *O*-dimensional hypercube with vertices defined by (±*t*, ±*t*,…, ±*t*). This occurs iff there exists a vertex of the hypercube and the origin lie on different sides of the hyperplane. Given a hyperplane defined by the equation *h*(***x***)) = 0 and a point ***x***_0_, *h*(***x***_0_) is proportional to the signed distance of ***x***_0_ from the hyperplane. Denoting a vertex of the hypercube as *t**x*** where ***x*** ∈{−1,+1}^*O*^, the above condition is equivalent to the product of signed distances of the origin and one of the vertices being negative, *i.e.*, there exists a ***x*** ∈ {−1,+1}^*O*^ such that

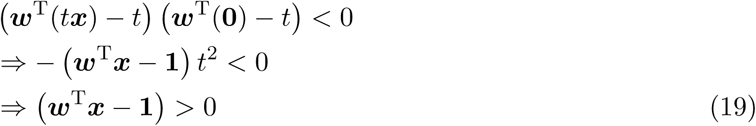

Equation 19 holds iff Σ_*j*_ |*w*_*j*_| > 1.

### 3 A comparison of the power of summary statistic imputation methods

Consider three SNPs, *snp*_1_*, snp*_2_*, snp*_3_ where SNPS *snp*_1_ and *snp*_2_ are observed while *snp*_3_ is unobserved. Assuming that SNP *snp*_3_ is the causal SNP with non-centrality parameter *λ*, the summary statistics at the three SNPs (*s*_1_*, s*_2_*, s*_3_)) follow the distribution:

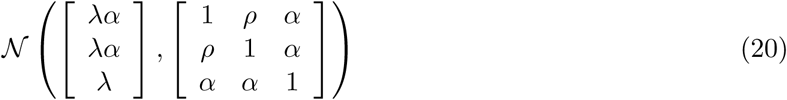

For this distribution to be well-defined, the covariance matrix must be positive-definite. A necessary condition for this is that the determinant of the covariance matrix (which is equal to the product of the eigenvalues) must be positive. Thus, we require the determinant (1−ρ)(1+*ρ−*2*α*^2^) > 0. |*ρ*| < 1 for the marginal distribution over SNPs 1 and 2 to represent a valid distribution. Further, we require 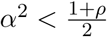.

Consider the case of SSI, *i.e.*, summary statistic imputation (with no variance re-weighting). In this case,

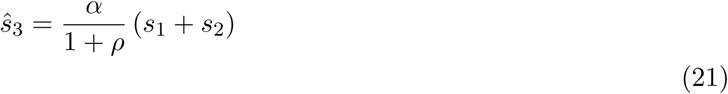

The mean, the variance and the coefficient of variation of 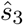 can be computed under the distribution (Equation 20):

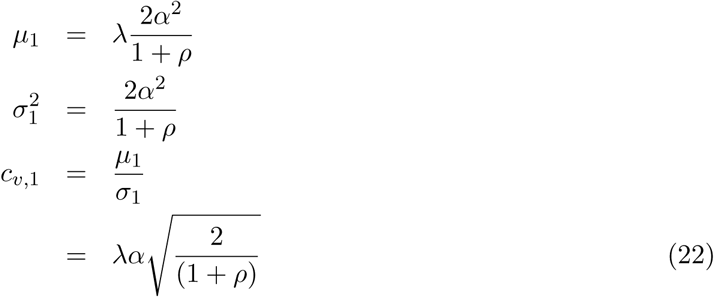

Now consider the case of SSI-VR, *i.e.*, summary statistic imputation (with variance re-weighting). In this case,

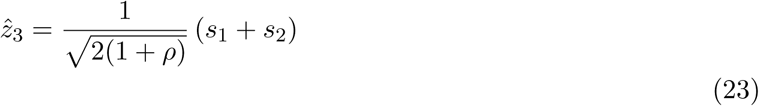

The mean, the variance and the coefficient of variation of 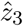 can be computed under the distribution (Equation 20):

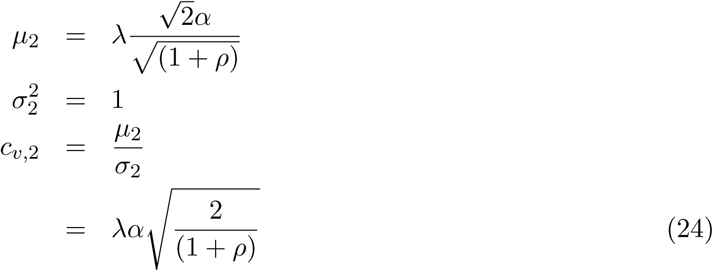

The power to reject the null hypothesis at a threshold *t* is

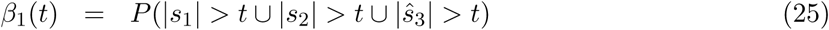

The power function depends on the joint distribution of *s*_1_, *s*_2_, and 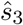. To simplify this expression, we will analyze the average power, a notion that is easier to analyze than the power as defined in Equation 25:

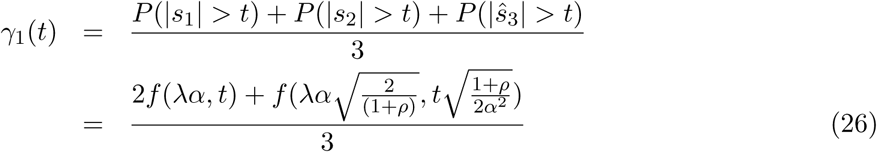

Here *f* is the power function defined in Section A.

Analogously, we can compute the average power *γ*_2_ for SSI-VR.

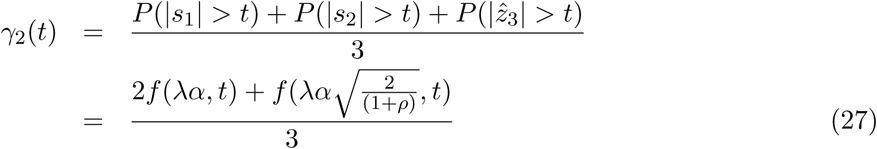

Comparing *γ*_1_(*t*) and *γ*_2_(*t*) allows us to understand the power of the two imputation methods. To analyze the power of each method, we need to do so at a threshold *t* such that each method attains the same FWER.

If *t*_1_ and *t*_2_ are the thresholds for SSI and SSI-VR, we have *t*_1_ ≤ *t*_2_. Thus, comparing Equations 26 and 27, we see that the first term for *γ*(*t*_1_) is greater than the first term for *γ*<u>(*t*_2_)</u>. The second term for *γ*(*t*_1_) will be greater than or equal to the second term for *γ*(*t*_2_) if 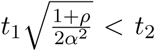. Thus, when the correlation between the unobserved and observed SNPs is close to its maximum possible value, *i.e.*, 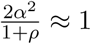, then SSI-VR has lower power than SSI.

#### A Power function

Given a normal random variable 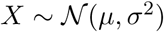, the probability that the absolute value of *X* exceeds a given threshold *t*:

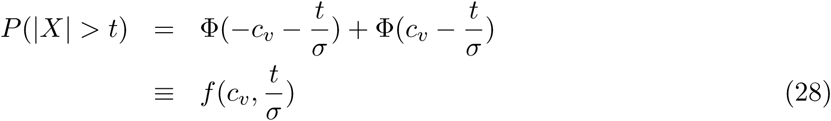

where 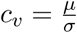 is the coefficient of variation and Φ is the normal CDF. Note that *f* increases with *c*_*v*_ and *σ* and decreases with *t*.

